# Genome Report: *De novo* genome assembly and annotation for the Taita white-eye (*Zosterops silvanus*)

**DOI:** 10.1101/2020.03.06.980599

**Authors:** Jan O. Engler, Yvonne Lawrie, Yannick Gansemans, Filip Van Nieuwerburgh, Alexander Suh, Luc Lens

## Abstract

The Taita White-eye (*Zosterops silvanus*) is an endangered songbird endemic to the Taita Hills of Southern Kenya, where it is confined to small areas of fragmented forest. With diversification rates exceeding those reported in most other vertebrates, White-eyes are a prime example of a ‘great speciator’. Nevertheless, we still know surprisingly little about the genomic underpinnings leading to this extraordinary fast radiation. Here, we present a draft genome assembly (ZSil_MB_1.0) for the Taita White-eye generated from a blood sample of a wild, female bird captured in the Taita Hills, Kenya. By performing a *de novo* assembly with linked-reads and annotation of the assembly with the MAKER pipeline, we generated a 1.069 Gb assembly with a scaffold N50 of 1.105 Mb and an L50 of 244. After quality evaluation of the assembly, we identified 92.1% of BUSCOs complete or fragmented, indicating that our *de novo* assembly is of high quality. This new assembly provides a genomic resource for future studies into the evolutionary and comparative genomics of this rapidly diversifying group of birds.

## Introduction

The Taita white-eye (*Zosterops silvanus*) is an East African highland songbird species endemic to the Taita Hills of Kenya, which has been subject to severe habitat loss over the past decades. The Taita White-eye is part of Zosteropidae, a unique family of birds known as “great speciators” (Diamond et al. 1976). Its most diverse genus *Zosterops* has one of fastest diversification rates among vertebrates, with over 100 species (and 250 sub-species; van Balen, 2019) emerging within the last 2.5 million years (Moyle et al. 2009). The phenotypic diversity in *Zosterops* is mostly conserved to a small body size and yellow-greenish colour patterns, even though aberrant phenotypes exist for some (mostly island endemic) taxa. With a lack of reference genomes available for this species and family as a whole, recent studies, such as on the silvereye (*Zosterops lateralis*, Cornetti et al. 2015) or the Mascarene grey white-eye (*Z. borbonicus*, Leroy et al. 2019) have demonstrated the relevance of generating genomic resources for this group of birds. Nevertheless, to further understand the rapid diversification in *Zosterops* more genomic resources from strategic phylogenetic positions would be highly important.

Here, we describe ZSil_MB_1.0, a new genome assembly using DNA extracted from a wild, female Taita White-eye collected from the Mbololo forest fragment in the Taita Hills, Kenya. To generate this assembly, we used 10X Genomics Chromium linked reads and the Supernova assembler, following by analysis of the assembly using the assemblathon2 Perl script (Bradnam et al. 2013) and annotation using the MAKER2 pipeline (v 2.31.10, Holt & Yandell 2011).

By generating the first de *novo* genome assembly for the Taita White-eye, we hope this will facilitate future research that aims to understand the underpinning of genotypic variation within this genus. In particular, identifying genomic regions under selection, which will greatly advance our understanding of the drivers of this radiation and help us to learn more about response to human activity and evolutionary constraints that might represent general features in other radiations too.

## Materials and Methods

### Genomic DNA Extraction and Genome Sequencing

We extracted genomic DNA from a blood sample of a female Taita White-eye captured on 15th July 1998 at the forest fragment of Mbololo in the Taita Hills, Kenya (ID: T29071_1006; GPS: 3°19’37”S, 38°27’05”E). DNA extraction was done with a standard phenol-chloroform protocol (Sambrook et al. 1989). A single 10X Genomics Chromium linked-read sequencing library was generated and sequenced on an Illumina HiSeq X instrument with 150-bp paired-end reads. In comparison to other short-read technologies, the Chromium system makes use of unique barcodes for each input DNA molecule which potentially allows for longer contigs and scaffolds of the assembly (Marks et al. 2019). Library preparation, sequencing and assembly – using the Supernova genome assembler for 10X Genomics (v. 1.1, Weisenfeld et al. 2017) – were performed at the National Genomics Infrastructure at SciLifeLab Stockholm, Sweden.

### Genome Assembly and Quality Check

We assessed the assembly quality using the assemblathon2 Perl script (*assemblathon_stats.pl*, Bradnam et al. 2015) slightly modified to be used in Strawberry Perl (64bit, v. 5.26.2.1) on a Windows 10 machine. Here, we include the following assembly statistics – for scaffolds and contigs respectively: the total number (N), number > 1000 bp, N50, L50, a well as maximum, mean, median, and total length. In addition, we used BUSCO v. 3.0.1 (Simao et al. 2015) on an Oracle VM VirtualBox for Windows (BUSCO_3_Ubuntu_Gnome_16.04 accessible through busco.ezlab.org) to assess the gene set completeness. In our case, we used the “odb9_aves” dataset to check our assembly for 4915 avian single-copy orthologs. BUSCO classifies orthologs as complete and single-copy (S), complete but duplicated (D), fragmented (F) or as missing (M). For comparison, we calculated the same statistics also for the silvereye assembly (Cornetti et al. 2015).

### Genome Annotation

Annotation of the white-eye genome was done using the MAKER2 pipeline (v2.31.10, Holt & Yandell 2011). Briefly, this software starts by masking repeat sequences using RepeatMasker (v4.0.7, Smit et al. 2017) with default settings. Next, MAKER2 uses protein sequences from related species to delineate homologous regions on the genome as candidate genes with blastx. We used protein sequences from chicken (*Gallus gallus*, NCBI GCF_000002315.4), turkey (*Meleagris gallopavo*, Ensembl UMD2), peregrine falcon (*Falco peregrinus*, NCBI GCF_000337955.1) and zebra finch (*Taeniopygia guttata*, Ensembl taeGut3.2.4). Then, the candidate gene sequences were retrieved from the genome and used to train a first *ab initio* gene predictor. The trained predictor was then used by MAKER2 for discovery of additional gene candidates. The resulting gene candidates were again extracted from the genome and used to train the same predictor for refinement or a different predictor for more specificity, which was then used by MAKER2 to extend and improve the set of identified genes. At each round of this iterative process, MAKER2 calculated a quality metric for each gene. The *ab initio* predictors used in this study were: SNAP (v2013-02-16, Korf 2004), AUGUSTUS (v3.2.3, Stanke et al. 2006) and GeneMark (v4.32, Ter-Hovhannisyan et al. 2008). Transfer RNA genes were identified using tRNAscan (v1.3.1, Lowe & Eddy 1997) in a final MAKER2 pass. Protein domain and GO term information was added using InterProScan (v5.29, Jones et al. 2014). Ribosomal RNA genes were identified with RNAmmer (v1.2, Lagesen et al. 2007). Putative functions were assigned to candidate protein coding genes using blastp and the complete Swissprot/Uniprot database. For the mitochondrial genome, we identified the relevant scaffold by comparison to the available silvereye mitogenome (Cornetti et al. 2015) and annotation was done using the MITOS software (v2.0.1, Bernt et al. 2013). Finally, all genome annotation data of *Z. silvanus* was bundled in a single GFF3 file.

### Data availability

Raw read data and the final assembly, ZSil_MB_1.0, are available from NCBI BioProject (PRJNA000000). Raw reads are available from NCBI SRA (XXX000000).

## Results and Discussion

Sequencing of the 10X Chromium library produced 370.81 million read pairs with a mean input molecule length of 38.6 kb and a median insert size of 320 bp, resulting in a 45x depth of coverage. The ZSil_MB_1.0 Supernova assembly produced 101,111 contigs with a total length of 1.017 Gb and a contig N50 length of 34.2 kb. The longest contig was 290.6 kb. These contigs were joined by Supernova into 55,959 scaffolds totaling 1.069 Gb in length and a scaffold N50 length of 1.105 Mb (Table 1). The longest scaffold was 8.001 Mb. For comparison, the silvereye assembly had a scaffold N50 length of 3.581 Mb and a contig N50 length of 41.7 kb (Cornetti et al. 2015), and the Mascarene grey white-eye had a scaffold N50 length of 1.763 Mb and a contig N50 length of 20.9 kb (Leroy et al. 2019). BUSCO scores indicated a high completeness with 92.1% of complete or fragmented single-copy orthologs and 7.1% missing orthologs for our Taita White-eye assembly (Table 2; compared to 96.1% / 2.9% for the silvereye and 97.3% / 2.5% for the Mascarene grey white-eye). The rate of duplicated orthologs was very low with 0.8% (1% for the silvereye). High BUSCO gene completeness has been found in most bird genomes (Peñalba et al. 2019) even though noticeable variation can be apparent among different assembly techniques and sequencing technologies (Peñalba et al. 2019, Peona et al. 2019). We therefore expect our BUSCO values to further improve in future assembly versions.

**Table 1:** Standard assembly metrics for scaffolds and contigs using the Supernova assembler including the total number (N), number longer than 1000 bp (N>1000), N50 and L50, as well as maximum, mean, median, and total length.

| Assembly statistic | Scaffolds | Contigs |
| --- | --- | --- |
| N | 55,959 | 101,111 |
| N>1000 | 55,920 | 88,537 |
| N50 | 1,104,592 | 34,226 |
| L50 | 244 | 8,377 |
| Max length | 8,001,144 | 290,647 |
| Mean length | 19,113 | 10,065 |
| Median length | 2,195 | 2,773 |
| Total length | 1,069,554,122 | 1,017,729,102 |

**Table 2:**
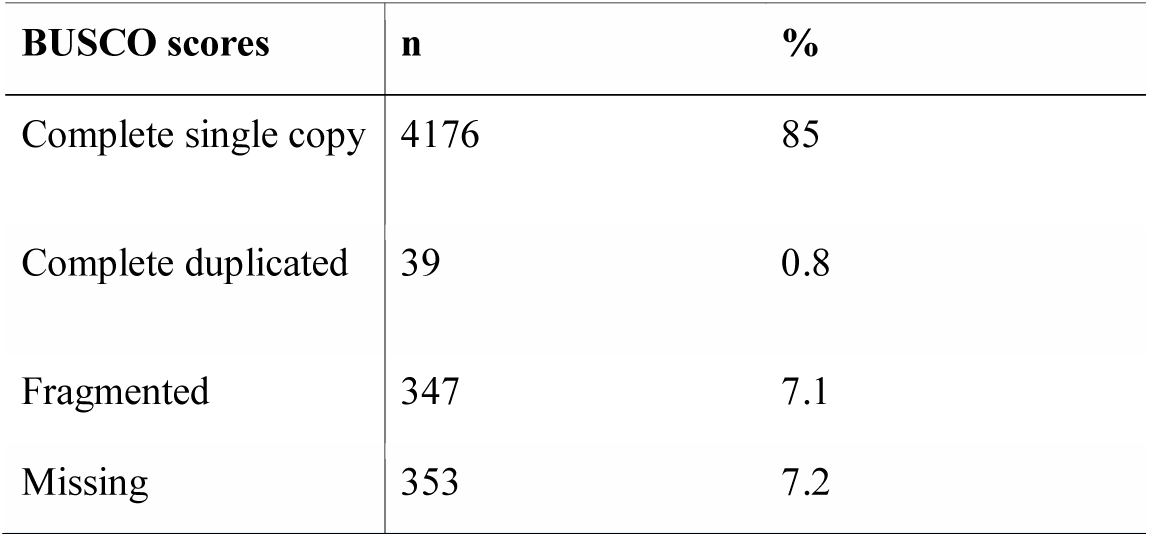
Assessment of the Supernova genome assembly and annotation completeness with BUSCO.

The final genome annotation using the MAKER2 pipeline revealed 18,033 candidate genes, of which 17,678 are potentially protein coding and of which 16,516 have been assigned with a putative function. Comparison of the protein-encoded genes having a known function with those extracted from the silvereye genome annotation (Cornetti et al. 2015) reveals 15,902 shared genes between the two species. Another 614 protein-coding genes with known function were only found in the Taita white-eye genome annotation, compared to 6,358 protein-coding genes solely found in the silvereye (Figure 1).

**Figure 1:**
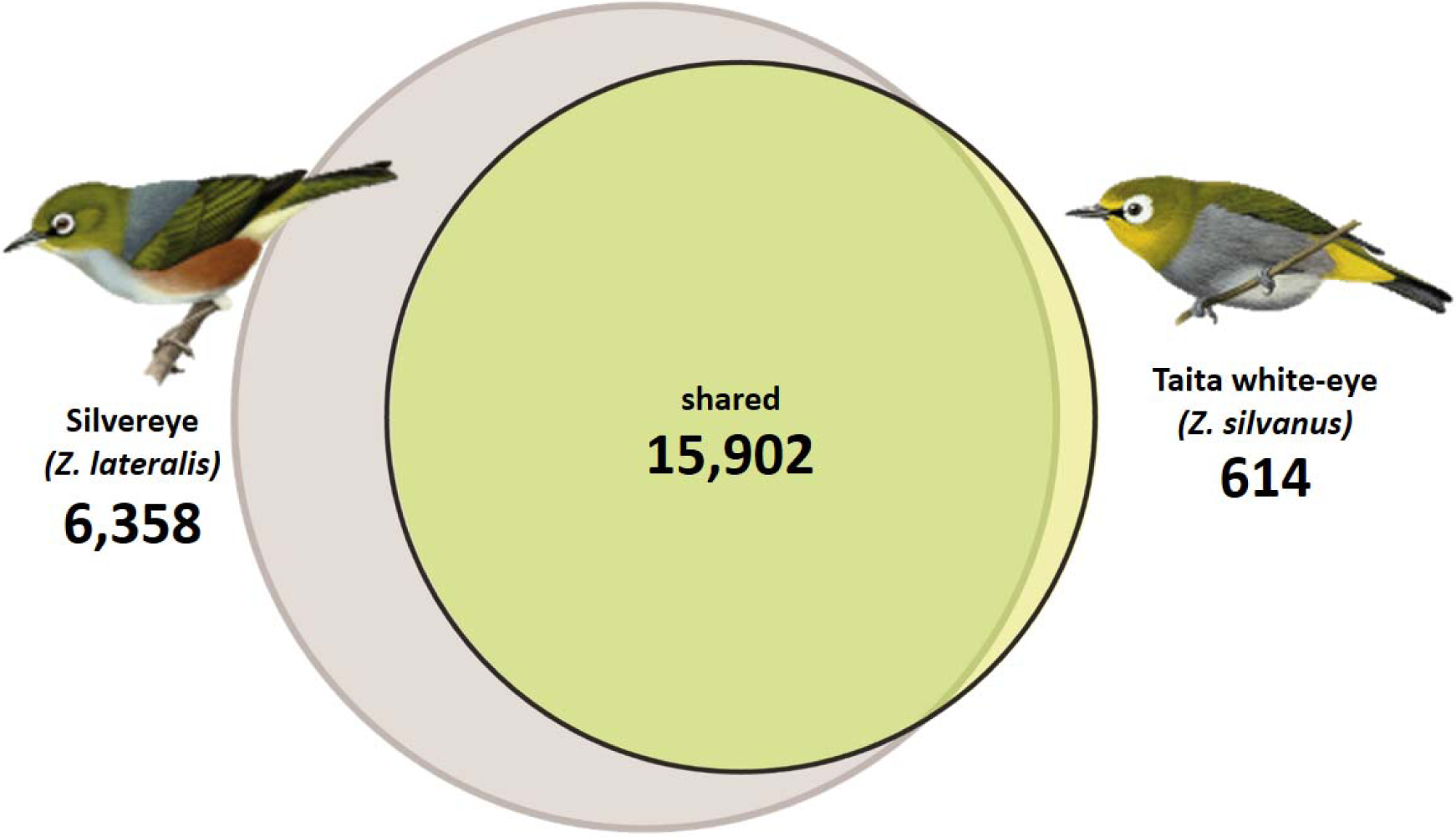
Venn diagram of predicted genes with putative protein function unique for as well as shared between the Taita white-eye (*Zosterops silvanus*, yellow circle) and the silvereye (*Z. lateralis*, grey circle) genome assemblies. Illustrations curtesy of HBWalive.

Linked-read sequencing has a huge potential for generating decent draft bird genome assemblies (Toomey et al. 2018, Boman et al. 2019, Kinsella et al. 2019, Lutgen et al. 2020). 10X Genomics Chromium is the most commonly used linked-read library preparation at present, but other linked-read methods (Chen et al. 2019, Redin et al. 2019) are equally promising but have yet to be tested on birds. Furthermore, the linked-read technology is still very new with ongoing developments and improvements of analytical tools and algorithms constantly being made. For instance, after the initial release of the Supernova assembly algorithm by 10X Genomics (v.1.1, Weisenfeld et al. 2017 – used in this report) there was a major update (v 2.x) after finalizing our analyses. In addition, further assemblers (e.g., ARCS, Yeo et al. 2017 cloudSPAdes, Tolstoganov et al. 2019, SLR-superscaffolder, Deng et al. 2019) and tools for downstream analyses (e.g. Tigmint, Jackman et al. 2018) recently emerged to improve such linked-read assemblies which might yield profound improvements of future versions of our draft assembly. Together in combination with other sequencing technologies such as Hi-C or long-reads (Peona et al. 2018), there is certainly room for further improvements towards a reference genome. Nevertheless, the current draft assembly ZSil_MB_1.0 marks an essential progress towards unraveling the genomic basis of diversification in a ‘great speciator’ system.

## Acknowledgments

The work was funded by the Research Foundation - Flanders (FWO; 1527918N & G042318N) to LL and JOE, and a SciLifeLab Swedish Biodiversity Program grant (2015-R14) to AS. JOE received additional funds by an FWO Postdoctoral Fellowship (12G4317N). The authors would like to thank Max Käller, Remi-André Olsen, and Joel Gruselius at SciLifeLab Stockholm, Sweden for conducting library preparation, sequencing, and assembly, as well as Viki Vandomme and Hans Matheve for managing the sample collection at the Terrestrial Ecology Unit, Ghent University. The authors acknowledge support from the National Genomics Infrastructure in Stockholm funded by Science for Life Laboratory, the Knut and Alice Wallenberg Foundation and the Swedish Research Council, and SNIC/Uppsala Multidisciplinary Center for Advanced Computational Science for assistance with massively parallel sequencing and access to the UPPMAX computational infrastructure. For comparison of the predicted white-eye proteins to those of the silvereye, we used its genome-derived peptide sequences and GFF3 file which were kindly supplied by Luca Cornetti.

JOE and LL conceptualized the research; YG and JOE performed analyses with critical input from FVN and AS; JOE and YL wrote the paper with critical feedback from all authors; JOE, AS, LL secured funding; all authors reviewed and approved the final manuscript.

